# Integrating case studies into graduate teaching assistant training to improve instruction in biology laboratory courses

**DOI:** 10.1101/594853

**Authors:** Jelena Kraft, Elise Walck-Shannon, Colleen Reilly, Ann E Stapleton

## Abstract

Today’s biology laboratory courses are primarily instructed by graduate teaching assistants (GTAs) working toward their own graduate degrees in a science field. Even though GTAs are the driving instructional force in biology laboratories, they receive little to no training in pedagogical principles and applications. Here, we present the GTAs’ self-reported unmet training needs with respect to teaching biology laboratory courses. To address these needs, we developed three training cases along with instructions for implementation in online and in-person GTA trainings. We piloted our training materials for GTA training in person at Iowa State University and in a Center for the Integration of Research, Teaching, and Learning three-part online workshop. Upon completion of these training sessions, GTAs reported an increase in both their confidence and knowledge of student-centered instructional approaches. Furthermore, examination of the online workshop participants’ lesson plans demonstrated that GTAs were able to implement effective teaching practices in their own lesson plans. For brief training sessions, these are positive outcomes suggesting these cases are effective tools that could be incorporated into a variety of GTA training programs to improve instruction in biology laboratory courses.

## INTRODUCTION

Given that a large portion of teaching, especially within introductory biology laboratories, is done by graduate teaching assistants (GTAs) (1, 2, 3), providing assistants with effective training has the potential to impact many students. In addition to GTAs filling the majority of instructional roles in laboratory courses, the introductory biology laboratories frequently enroll large numbers of students (2) such that GTAs have an impact on a multitude of undergraduates in science, technology, engineering, and math (STEM) disciplines – which further highlights the need for rigorous GTA training programs. Many of these GTAs will eventually become faculty members, making training an important investment for tomorrow’s students as well. Despite the central role GTAs play in student learning in these disciplines, GTA training programs often fall short in providing pedagogical training outside of the GTAs’ specific scientific disciplines (3).

The format and content coverage of GTA trainings varies widely (3, 4, 5), but training rarely occurs concurrently with the course that GTAs are teaching. The most frequent type of GTA training includes one-time training at the beginning of the semester, typically limited to a few hours (5). This type of training has limited benefits as GTAs are less likely to transfer the concepts and practices they learn into their own teaching over the long term (3, 6, 7).

In addition to temporal limitations, typical GTA training programs are hindered by the narrow scope of discussed topics. GTA trainings frequently cover topics related to general laboratory policies and course management and fail to address theories about how students learn or cover evidence-based teaching practices that would be the most beneficial to biology lab GTAs and their students (5, 6, 8). Course, laboratory, and institutional policies and discipline-specific knowledge are essential components of laboratory instruction; however, unless GTA trainings also address pedagogical strategies, they are unlikely to translate into high-quality science instruction.

Our goal was to identify any unmet training needs of GTAs in biology labs and provide them with just-in-time materials to help prepare them for lab-specific challenges. Towards this goal, we surveyed GTAs across three universities to elicit what they perceived to be the greatest challenges to effective teaching in biology lab courses. From this survey, we identified the need for additional training in determining their own teaching style, managing time efficiently, and motivating students to learn. We addressed these unmet training needs through the development of training case studies designed specifically for biology labs and used the cases for GTA training via in person and online workshops. Because engaging in the deliberate practice of teaching techniques during training is essential in order to prompt the graduate lab instructor to use those techniques in their own classrooms (7), we incorporated construction of lesson plans and peer and instructor feedback into our online training. We also modeled an active learning process in our online workshop design, as many lab instructors reported no prior experience with active learning classroom methods. These types of methods have been shown to improve students’ learning outcomes in a variety of classroom settings in STEM disciplines (9, 10). The effectiveness of GTA trainings using the case-study based training materials were assessed using the Knowledge Survey (KS) (11, 12), the Center for the Integration of Research, Teaching, and Learning (CIRTL) post-workshop survey, and the analysis of the workshop participants’ lesson plans. Below we offer recommendations about how to integrate these case studies into future GTA training.

## METHODS

### Design, administration, and analysis of GTA training needs survey

To identify the problem areas for GTAs working in biology labs, two faculty members involved in the undergraduate biology education at Iowa State University (J.K.) and University of North Carolina Wilmington (A.S.) developed a training needs survey. The survey contained 2 Likert-type (Appendix 1, questions, categories, and frequency responses) and 4 open-ended questions (Appendix 2, questions, categories, and frequency responses). The questions probed into the following three categories of GTA teaching experience: 1) demographic information (Supplemental Table A1), 2) personal reflections on teaching biology labs, and 3) attitudes toward teaching. The survey was administered in the Spring 2017 semester with 137 invites sent by email to GTAs working in biology labs at Iowa State University, University of North Carolina Wilmington, and University of Georgia. We received 49 GTA responses: 35 from Iowa State University, 2 from University of North Carolina Wilmington, and 12 from the University of Georgia. The frequency of responses to Likert-type questions were recorded for each answer choice category using Qualtrics software, Version (January-February 2019). For the open-ended survey questions, J.K. and A.S. read, categorized, and counted GTA responses for each question reported here (Appendix 2, questions, coding, and frequencies for responses). Surveys and research protocols were approved by the Institutional Review Board (**IRB ID 17-104**) at Iowa State University.

### Training case studies writing

After analysis of the survey data, we developed three training cases to demonstrate to GTAs how to address one of three significant pedagogical goals: 1) incorporate active learning methods to break up lectures, 2) effectively assess student learning of lab material, and 3) incorporate real-world examples in lab lesson plans to increase students’ motivation. Each case presented a scenario in which a GTA teaching a basic biology laboratory concept is challenged to address one of these three pedagogical goals. A set of discussion questions accompanied each case, inviting GTAs to work together to apply the discussed teaching theory and methods and then propose some solutions to overcome the instructional hurdle. Finally, GTA discussion of each case concluded with an application exercise, where the lab lesson outcomes were outlined and GTAs were charged with developing one component of a lesson plan (e.g. a brief lesson introduction, formative and summative assessment, or an active learning strategy) to integrate the discussion and theory into practice (Appendix 3, training cases and accompanying activities full details).

### Supporting training materials

In addition to the three training cases for GTA trainings, we adapted a lesson plan guide (13, 14) using the backward design pedagogy (15) to specifically demonstrate how the developed cases fit into lesson planning (Appendix 4). This lesson plan guide was used as a central organizing theme for GTA trainings, in which learning outcomes were explained using Bloom’s Taxonomy (16) and student-centered instructional approaches and assessments were explained using constructivist learning theory (17). The training cases, discussion questions, application exercises, and lesson plan guides were all integrated into interactive PowerPoint presentations to illustrate the example pedagogy during workshop delivery. Additional training materials used for GTA training in an online workshop can be accessed at https://www.cirtl.net/resources/700.

For the delivery of the workshop trainings, we adapted the cases and materials to the online, synchronous setting while accounting for CIRTL guidelines and Moodle platform details. To compare the in-person workshop instructor materials to the online workshop materials we calculated semantic similarity by input into the online comparison tool at *http://swoogle.umbc.edu* (18).

### Delivery of training

Training materials were used for GTA training in person at Iowa State University and in the Center for the Integration of Research, Teaching and Learning (CIRTL) three-part online workshop titled “Integrating Effective Teaching and Assessment Practices in Biology Lab Courses through Case Studies” during the Spring 2018 semester.

In-person training was integrated into the pre-existing mandatory pre-semester orientation session for GTAs working in the principles of genetics laboratory (an upper division course for majors). Five GTAs, who were working towards their PhD degrees in a related field and assigned to teach two sections of this lab in the Spring 2018 semester, participated in this training (Table 3). The training lasted four hours: the first two hours of training included general lab and safety orientation, while the remaining two hours were dedicated to the pedagogy related to effective teaching and assessment practices in biology lab courses and incorporated the three twenty-minute training case study discussions. A single instructor (J.K.) led the training and facilitated the case discussions. No homework was assigned to this group.

Online training consisted of three workshops, which were presented weekly. The first two workshops were seventy-five minutes long, during which GTAs received instruction in effective teaching and assessment practices and worked in groups on the case study discussions and lesson planning. Each group contained six randomly assigned workshop participants, though not all were active participants (Table 3). After each session, participants were assigned homework consisting of continuous development of a lesson plan on a biology lab topic of the participant’s choice. This homework incorporated the pedagogy discussed in the case studies that were presented in the first two workshop sessions. After each session, participants received a review from at least one other peer and a few participants requested additional feedback from the instructors. The final session included a 90-minute delivery and discussion of participants’ lesson plans with simultaneous feedback from peers and instructors. Three instructors/moderators directly facilitated the workshop training (J.K., A.S. and E.W.S.) and one additional moderator (C.R.) provided online teaching expertise to the moderators and monitored/provided comments to the participants in the chat window. Workshop facilitators (J.K., A.S., E.W.S., and C.R.) completed the mandatory CIRTL instructor training and developed the workshop facilitator training delivery plan that followed the trainings.

### Assessment of training

The impacts of these cases on GTAs’ understanding of different pedagogical approaches to effective lab instruction was evaluated via the Knowledge Survey (11, 12) for both the in-person and online training group before and after training. The survey was formatted for online delivery using Qualtrics software, Version (January-February 2019) (Appendix 5).

#### MANOVA and descriptive statistics for pre-post survey analysis

The KS pre- and post-survey data were formatted for statistical analysis (Supplemental Maerials File titled KnowledgeSurveyMaster.xlsx). We used JMPv13 (SAS, Inc, Cary, NC) to fit distributions to the pre- and post-survey data to select the best-fitting distribution for repeated measures ANOVA. We fit the model using the normal distribution with survey question nested within individual (for more statistical model fit detail, please see Appendix 6 Data Analysis Model Fit Details).

Additionally, for the online workshop training group, we evaluated participants’ ability to incorporate teaching practices by analyzing GTAs’ lesson plans and using the CIRTL standard post-survey (Appendix 7, CIRTL post-survey results, and frequency responses). Workshop discussions and polls were used to characterize participants’ experiences with developing lesson plans (Appendix 8, Analysis of in-workshop live polls and discussions).

## RESULTS

### Unmet training needs

To characterize the unmet training needs for the biology laboratory GTAs, we analyzed the training needs survey results from the survey that was administered to GTAs at Iowa State University, University of North Carolina Wilmington, and University of Georgia in the Spring of 2017. The questions on this survey included those relating to their current experiences with teaching as well as their attitudes toward teaching. The demographics of GTAs participating in the survey are listed in Table 1.

**Table 1:** Characteristics of GTA who completed the training needs survey. GTA: Graduate Teaching Assistants

| Institutions | Lab courses taught | Semesters as a lab TA | Lecture courses taught | Semesters as a lecture TA |
| --- | --- | --- | --- | --- |
| Iowa State University | Biochemistry | Zero: 2.22% | Introductory Biology | Zero: 69.44% |
| University of Georgia | Botany | One: 26.67% | Genetics | One: 19.44% |
| University of North Carolina Wilmington | Developmental Biology | Two: 20.0% | Cell Biology | Two: 8.33% |
|  | Ecology | Three: 8.89% | Botany | Three: 2.78% |
|  | Genetics | Four and more: 42.22% | Biochemistry | Four and more: 0.0% |
|  | Human Anatomy and Physiology |  | Biophysics |  |
|  | Introductory Biology |  |  |  |
|  | Molecular Biology |  |  |  |
|  | Organismal Biology |  |  |  |

Most GTAs had at least one or two TA lab experiences and the majority were experienced TAs, having supervised four or more labs. Most of these GTAs (70%) had not served as a TA in a lecture class. The labs they taught ranged from introductory biology/biochemistry to specialized lab topics in animal and plant systems. GTAs indicated that they were interested in teaching prior to assuming their positions and felt confident in their ability to do so, but were not overly confident in teaching the lab they were assigned (Table A1 in Appendix 1). On this survey, GTAs also agreed that lecturing is still a primary way of conveying content in the lab. The majority of GTAs expressed interest in learning to use more active teaching methods in their lab and believed that this would enhance their teaching ability and student learning. GTAs who took the survey viewed their work positively and would recommend these position to their peers (Table A1, Appendix 1).

In one of the open-ended questions on this survey, we asked GTAs to tell us about what they wished someone had told them about teaching a lab class. Their answers fell into four categories: 1) how to come up with their own teaching style/teaching strategies that work (39% of responses), 2) how to manage time efficiently (29% of responses), 3) how to motivate students to learn (15% of responses), and 4) how to deal with difficult student/situations in the lab (17% of responses) (Figure 1).

**Figure 1:**
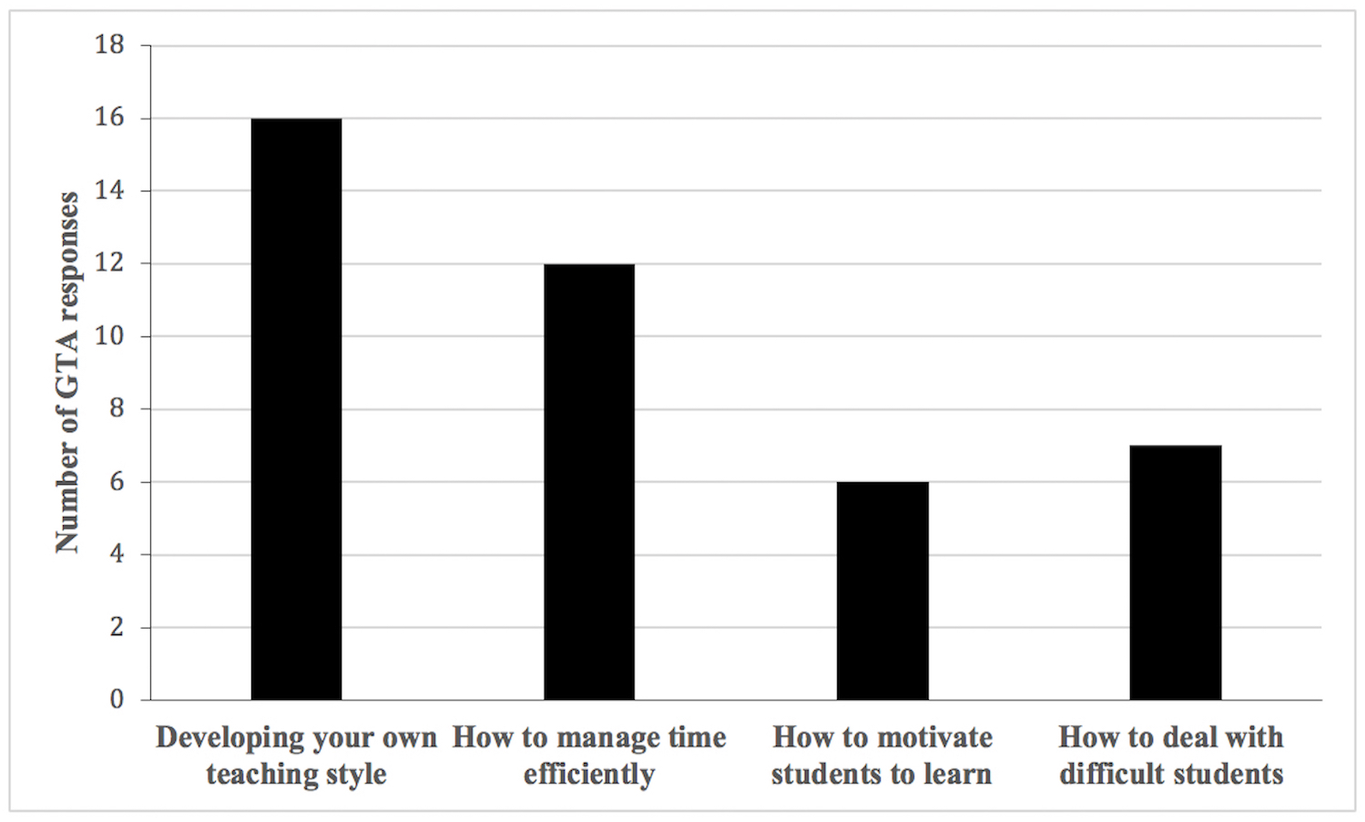
GTA responses (N=41) on the training needs survey to the open-ended question about “What do you wish someone had told you about teaching lab courses?” Some GTA answers fit more than one category and not all GTA provided an answer or detail relevant to the question asked.

Coincidentally, in the follow-up question on this survey, GTAs were asked about the single most important thing they would tell a colleague about being a TA, and the responses matched the four categories listed above (Figure A2.1 in Appendix 2). Additional open-ended questions on this survey that probed into GTAs’ experiences with teaching a biology lab course were consistent with instruction, grading, student motivation, and classroom management being the most most labor intensive parts of TAing a laboratory course (Figure A2.2, in Appendix 2) and were something GTAs at the three institutions wished they had more time to adequately prepare for (Figure A2.3, in Appendix 2).

Based on these results, we proceeded to develop three training cases (full case details are available in Appendix 3) around implementing active learning methods, effectively assessing student learning of lab material, and improving student motivation by incorporating real-world examples into lesson plans. Each case outlined a lab scenario in which a GTA is depicted teaching a general biology lab concept and encounters some type of student resistance to learning the material. Each case is followed by questions that probe the instructional deficiencies presented in the case and activities that engage GTAs in applying the pedagogical practices discussed. We interwove some commonly encountered classroom management situations, such as disinterest or confrontation about learning a lab concept, poor performance on the assessments, and technology distractions, into the three cases we developed.

### In-person training

Developed cases were used for the training of GTAs in person in January of 2018. Five GTAs at Iowa State University attended the in-person training session. The training session was incorporated into an existing pre-semester orientation training session in order to get GTA buy-in. The last two hours of the pre-semester orientation were dedicated solely to case study readings and group work discussions.

The GTAs in this group either had no previous teaching experience (two out of five) or were experienced GTAs (three out of five), and all five GTAs were assigned to teach an introductory genetics laboratory (Table 2).

**Table 2:** Prior teaching experience of GTA who participated in trainings and completed the knowledge pre- and post-surveys

| Training type | Number of GTA participants | Zero semesters | One semester | Two semesters | Three semesters | Four or more semesters | Laboratory course taught |
| --- | --- | --- | --- | --- | --- | --- | --- |
| In person | 5 | 2 | 0 | 0 | 0 | 3 | Developmental biology<br>Genetics<br>Introductory biology |
| Online workshop | 13 | 4 | 1 | 1 | 1 | 6 | Developmental biology<br>Environmental biology<br>Genetics<br>Introductory biology<br>Introductory geology<br>Organismal biology |

In this group, GTAs were first introduced to the pedagogy related to the first two steps of a lesson plan (Appendix 4, lesson plan guide) followed by the case study discussions on “Motivating students through real-world research examples” (Appendix 3, training cases). Next, the GTAs were introduced to the pedagogy of steps three and four of the lesson plan (Appendix 4, lesson plan guide) followed by the case study discussions on “Active learning method” and then “Effective assessment of biology lab work” (Appendix 3, training cases). Due to time constraints, GTAs in this group did not work on developing their own lesson plans.

### Online workshop training

In addition to in-person training, the cases were used as a framework for training GTAs in an online format as part of a Center for the Integration of Research, Teaching, and Learning (CIRTL) workshop titled “Integrating Effective Teaching and Assessment Practices in Biology Lab Courses through Case Studies” in Spring 2018. Thirty-five participants across eleven institutions in the CIRTL network were enrolled in this workshop. Of these, only thirteen participants completed both the KS and CIRTL workshop surveys indicating their previous teaching experience (Table 2). As in the in-person training group, most GTAs participating in this workshop either had no experience or were highly experienced TAs. There was a greater range of previous lab teaching experiences for the CIRTL workshop group compared to the in-person training group.

The workshop was presented in two seventy five-minute long sessions and one ninety-minute long session. In workshop one, participants were first introduced to the general lesson planning steps, focusing on formulating learning outcomes, and developing a lesson introduction as was done in the in-person training group. This brief instruction was followed by small-group discussions about the “Motivating students through real-world research examples” case study; participants then worked in groups to generate a lesson plan template with learning outcomes and an introduction for the biology lab topic presented in the case study. Upon conclusion of the first workshop, participants were asked to use the content discussed to develop a working draft of a lesson plan focused on a biology lab topic of their choice and submit it for peer review.

In the second workshop, participants learned about different forms of active learning and assessment followed by a group discussion focused on the case studies “Active learning method” and “Effective assessment of biology lab work.” Following the discussions, participants continued to work in groups on a lesson plan template that incorporated at least one type of formative and summative assessment and one active learning approach. At the end of workshop two, participants were asked to return to their individual lesson plans, develop steps three and four, (Appendix 4, lesson plan guide), and then to submit it for additional peer feedback.

The final workshop session focused on small group presentations of the participants’ lesson plans. GTAs were also asked to include strategies for facilitating student learning during lab and a reflection about teaching (steps 5 and 6 of lesson plan guide, Appendix 4), although we did not offer explicit instruction beyond suggested readings on the topic. Each presentation was followed by synchronous peer and instructor feedback.

### Evaluation of GTA trainings

Following the trainings, the Knowledge Survey (11, 12) was administered to measure any shift in GTAs’ perceived understanding of the pedagogies related to the use of assessments, explanations, and active learning to promote student learning. On this survey, GTAs in both training groups reported their confidence in knowledge of items across the first five learning goals (31 items) on a scale of 1 to 3, with 1 indicating lowest confidence and 3 indicating most confidence (Appendix 5, KS Qualtrics survey).

Comparison of the Pre- and Post-KS scores for the two training groups indicated that both groups made a shift towards an increase in confidence in comprehension of the teaching items on the KS survey (Figure 2).

**Figure 2:**
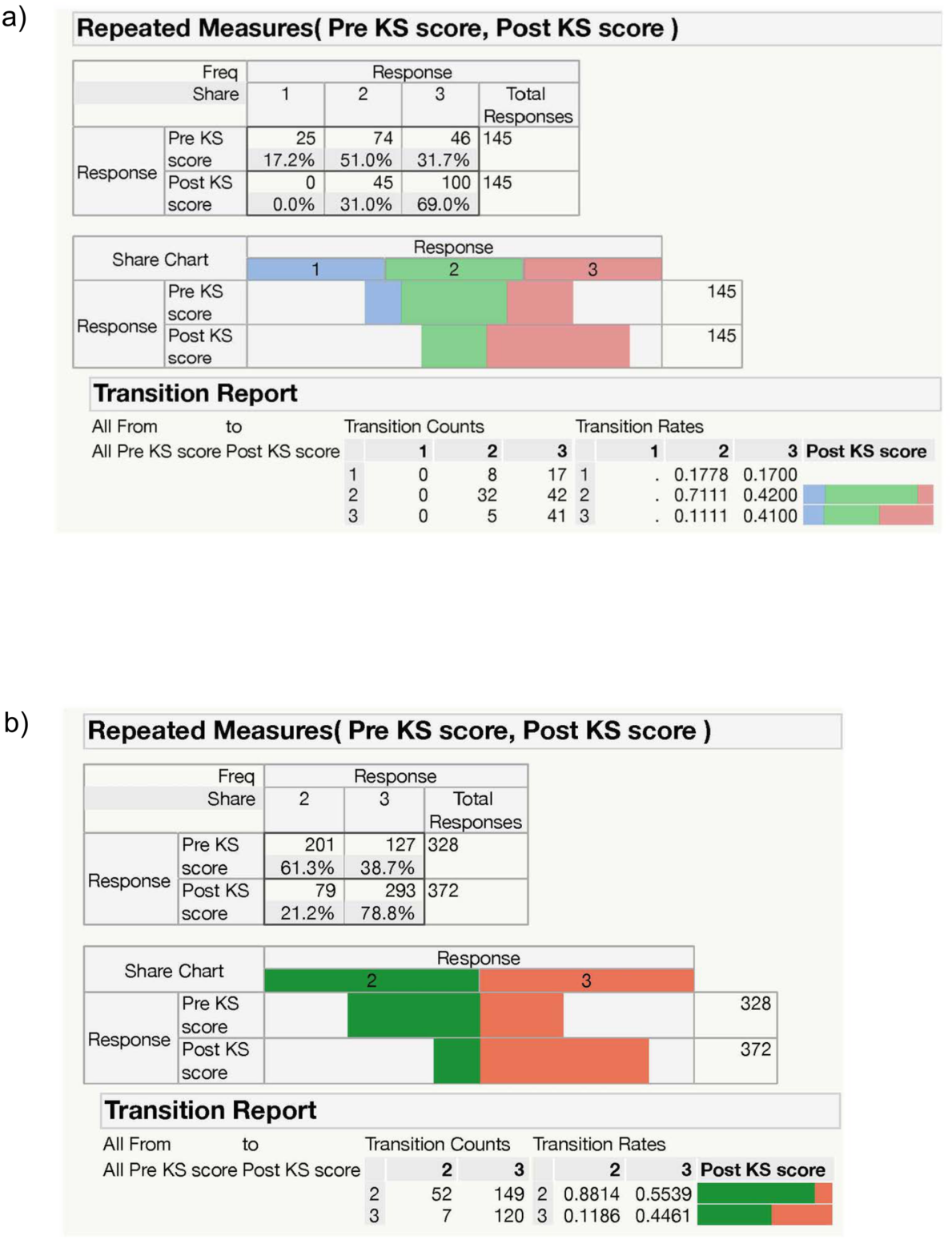
Shift in survey scores pre- and post-KS across all survey questions and individuals. a) in person b) online workshop. GTAs reported their confidence in items relating to these learning goals on a scale of 1 to 3, with 1 indicating lowest confidence and 3 indicating highest confidence. Both training groups completed the pre-KS before the training and post-KS upon completion of the training. KS questions and the scoring rubric were identical on the pre- and post-survey administration.

The MANOVA analysis indicated a significant difference between pre- and post- (P<0.0001 for in-person and P<0.0001 for online), with no significant difference between survey questions. The survey numbers are rarely lower in the post-analysis, though some GTAs start with high confidence and remain highly confident; in the online workshop, 3 of the 13 individuals had one or two questions with lower values in the post-survey (grey background in multi-box figure in Appendix 6 Figure A6.2).

GTAs reported their confidence in items relating to these learning goals on a scale of 1 to 3, with 1 indicating lowest confidence and 3 indicating highest confidence. Both training groups completed the pre-KS before the training and post-KS upon completion of the training. KS questions and the scoring rubric were identical on the pre- and post-survey administration.

For GTAs who were trained in-person across five learning goals, the distribution of responses shifted from 17.2% lowest, 51% moderate, 31.7% highest confidence on the pre-KS to 31% moderate and 69% highest confidence on the post-KS (Figure 2a). In the online workshop group, there were no GTAs who lacked knowledge of the items on the pre- or post-KS survey (Figure 2b). For the online workshop group, the confidence scores shifted from 61.3% moderately and 38.7% highly confident on the pre-survey to 21.2 % moderately to 78.8% highly confident on the post survey (Figure 2b). In terms of the transition rates, the in-person training group had 8 responses that shifted to moderately confident or 17 to highly confident pre- to post-KS (Figure 2a). In the online workshop group, there were 149 responses that transitioned to highest confidence pre-to post-KS (Figure 2b).

For the CIRTL workshop participants, we were also able to examine the participants’ ability to incorporate the concepts into their lesson plans. The majority of workshop participants submitted a lesson plan, and we found that each one included a learning objective(s), at least some mention of active learning techniques, and some plan for assessment. Additionally, each active participant revised that plan once after receiving feedback from their peer(s). In the final session, participants also received feedback from one of the course facilitators (though this feedback was rather brief in this online workshop setting). These results are supported by the KS pre-/post-survey, which found that participants had gains in their perceived ability to “use principles of learner-centered teaching to construct a lesson plan.” At pre-survey, 11.1% said that they felt confident that they could do this currently; whereas, at post-85.7% felt this way (Table 3).

**Table 3:**
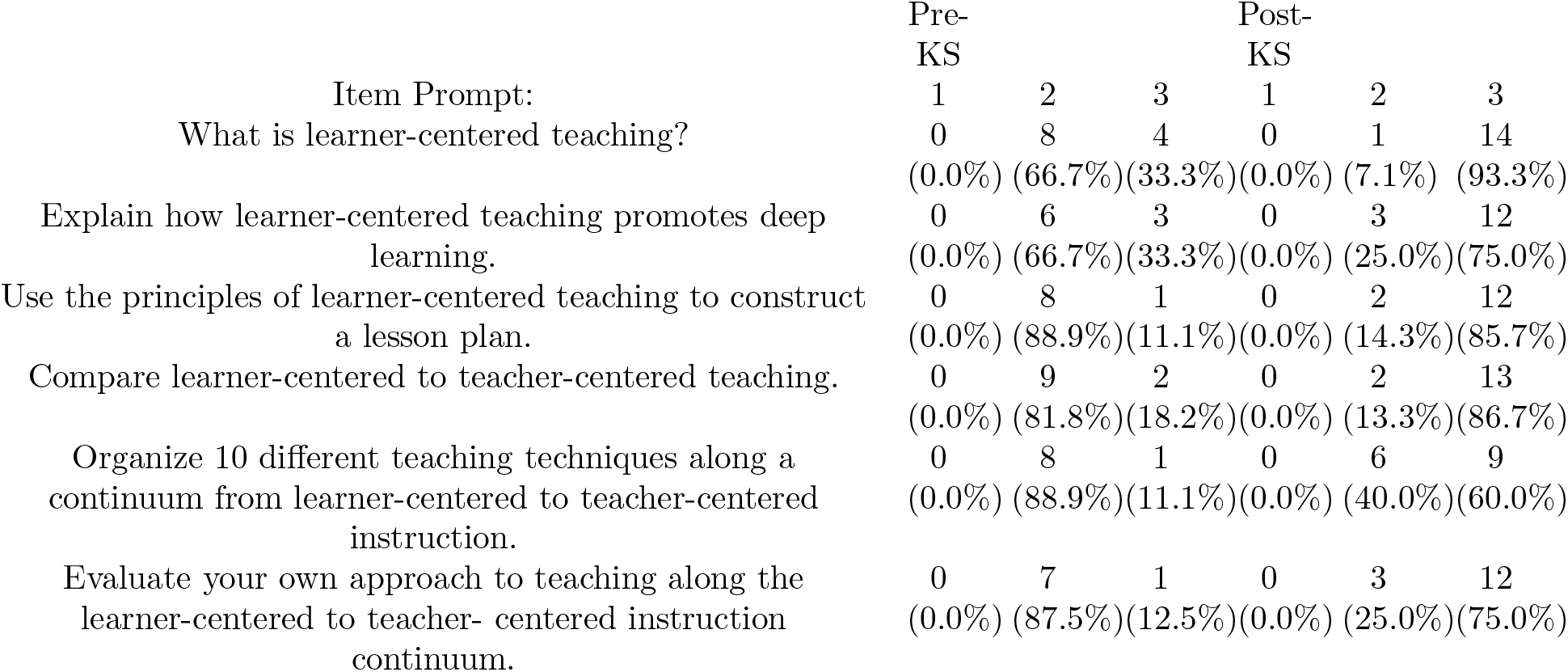
Partial item-by-item data for online workshop participants taking the KS. Scale for responses: 1 = I do not understand the question, I am not familiar with the terminology, or I am not confident that I can answer the question well enough for grading purposes at this time. 2 = I understand the question and: I am confident that I could answer at least 50% of it correctly, or I know precisely where to find the necessary information and could provide an answer for grading in less than 20 minutes. 3 = I am confident that I can answer the question sufficiently well-enough for grading at this time.

Additionally, during the final presentations, we observed that one participant developed a novel case study on a biology lab topic and incorporated it into their lesson plan as a form of student-centered group activity. For these reasons, we conclude that the majority of active participants gained both a knowledge of pedagogy and the skill set necessary to apply it to their teaching. Acquiring this skill set is more likely to translate into continued advances in teaching practice.

As part of the participants’ presentation of their lesson plans, we queried the participants about the strategies they employed to develop their lesson plans for a biology lab of their choice. For example, one of the CIRTL workshop participants described their process as follows: “So when I’m planning a lesson the first thing that I often do is to think about what I want students to *gain* from that lesson, and then I usually use my goal as my starting point and go from there.” This participant was not an exception; we observed other workshop participants being highly engaged with the cases during group discussions, contributing ample ideas in response to these specific cases. In contrast, other GTAs initially struggled to apply the concepts from the cases (i.e., incorporating real-world examples) to their own lessons.

One week after the initial workshop, participants were still reporting difficulty in applying their real-world introductions to their own lessons. This struggle was apparent when participants were asked what was “the most difficult part for you in preparing [your] lesson plan” at the start of the second session. The majority (66.6%, 8/12) mentioned the real-world introductions, a topic directly covered in the first case study (Table 1 of Appendix 8, “live” CIRTL workshop data analysis). Their concerns were split between wanting to choose an introduction that was interesting to students (N=3), developing an introduction that was at the proper level of detail for students (N=3), and developing an introduction that took up an appropriate amount of time (N=2). The remaining participants felt difficulty in generating learning outcomes, which were also covered in the previous session. Together, this suggests that incorporating real-world examples to inspire students in a laboratory introduction is a difficult skill to master. For this reason, it is appropriate to focus on this in training sessions, especially when constructive discourse—generating and communicating ideas (i.e., presenting lesson plans) and receiving and responding to feedback (i.e., revising lesson plans from peer review comments)—is incorporated.

Following the last workshop session, participants were instructed to complete the standard CIRTL-post survey. On this survey, all of the workshop participants responded that they were either likely or very likely to integrate what they have learned during the workshop into their research, teaching, or work environments (Figure 7.1, Appendix 7). The overwhelming majority of participants agreed or strongly agreed that the workshop training will help them prepare for a teaching career. They also indicated that workshop discussions were valuable for the development of their own lesson plans (Table 7.1 of Appendix 7, CIRTL Post-Survey questions and analysis). A number of our participants indicated that they benefited from discussing the case studies during our workshop sessions; however, a few identified this as a weak part of the workshop experience (Table 7.2 of Appendix 7, CIRTL Post-Survey questions and analysis). Representative examples of participant’s comments about usefulness of cases for training include the following:

> “I enjoyed discussions and the case studies. It gave me a chance to hear answers that were different from my own and from a perspective I may not have considered.” “I felt most engaged during the small groups discussing case studies. The size of the group, the lack of leader/organizer, the assigned roles, and the questions posed worked well for encouraging discussion.” “I felt the most engaged when discussing case studies with my breakout group and during the final presentations of lesson plans. In the former case, I already knew much of the material being presented by the moderators, so discussions of how to apply the knowledge to a lab setting were the only new and interesting parts for me. In the latter, I really enjoy providing feedback and was excited to see more of the diversity of ideas developed by the larger group than just my two breakout group partners.”

For a brief, two-hour long in-person training and three-part workshop, these are encouraging outcomes. Our evaluations indicate that the targeted training cases can be a valuable tool to focus participants’ attention on needed skills (i.e., framing lab topic, grading efficiently, and active learning) and provide strategies for using those skills in the creation of their own lesson plans.

## DISCUSSION

While training programs for biology GTAs are now a common practice across the country, they are still limited in duration and breadth (5). Here, we surveyed GTAs across three graduate biology institutions to identify specific areas of focus for GTAs’ training in preparation for teaching a biology lab course. The respondents reported a need for instruction in determining their own teaching style, crafting workable teaching strategies, managing time efficiently, and motivating students to learn. These responses are also consistent with the last national survey of GTA professional development practices in which these areas of training were most requested and also least likely to be taught during GTA trainings (5).

Towards this goal, we developed three training case studies that incorporated discussion and group work and, in the case of the online training group, collaborative and an individual work on lesson planning. For the majority of GTAs, we saw increase in confidence pre- to post-trainings in the in-person and online training groups as indicated by the KS survey. However, we observed that a few online participants had a decrease in confidence, which can be an important part of learning as students transition from a state of unconscious incompetence to conscious incompetence (see 19 for an explanation of the conscious-competence model). Also, evaluation of at least one other training program (4) has shown a misalignment between the quality of artifacts versus self-reported confidence (better alignment and student-centered approaches but less confidence).

Increases in GTAs’ confidence in the knowledge of evidence-based instructional practices pre- to post-training, while encouraging, are unlikely to immediately translate into teaching practice. Experiences with similar training programs demonstrate that even when discipline-specific pedagogy is explicitly covered in trainings, GTAs are unlikely to stably integrate discussed practice into their teaching unless they are given ample opportunities for practice and feedback (6, 7, 20, 21). We noticed that there were participants in our online training group who, despite being given an opportunity to practice lesson planning steps with their peers during workshop and then to apply it on their own lesson plan, still struggled to do so after the first two workshops. Ongoing work and feedback on lesson planning was beneficial to all active participants, and they all presented fully developed lesson plans (on steps 1-4 of lesson plan, Appendix 4) by the last workshop. However, we note that the components of lesson planning that were mentioned but not included in direct instruction, such as steps on how to facilitate student learning and reflections about teaching (Steps 5-6 of lesson plan, Appendix 4), were missing in all of the participants’ lesson plans. For our GTAs, both targeted pedagogical training connected to the science discipline and the opportunities for practice during and after the trainings were more likely to result in GTAs’ adoption of evidence-based teaching practices into their lesson planning. While these observations (that continued practice is more beneficial for learners’ development) are not new, it highlights the importance of ongoing discipline-specific pedagogical training for GTAs to become confident and purposeful about implementing student-centered teaching approaches into their teaching.

### Limitations and recommendations

We note that one of the study limitations was the small sample size, especially for the in-person training group which consisted of only five GTAs. However, the sample size is on par with similar studies on biology GTA professional development (for example, 4, 11, 22, 23, 24, 25). To leverage the small sample size, we focused how GTA’s confidence changes from pre- to post-trainings and, qualitatively, on the demonstration of an ability to incorporate pedagogy into lesson planning by the online training group participants.

Another limitation is the short duration of training for both GTA groups, but especially for the in-person training group. For this group, we were only able to ask about GTAs’ confidence in knowledge of pedagogical approaches covered in the training and did not actually observe their resulting teaching practices. Additionally, while we were able to observe if and how participants in the online workshop were incorporating pedagogy into their lesson plans, we do not have any data on how many of the GTAs incorporated the pedagogical strategies they learned into their teaching at their home institutions. Comparison of the in-person workshop instructor materials to the online workshop materials using semantic similarity resulted in a score of 81% similar at the phrase level, so the differences in training material could also contribute to the different patterns observed in these two settings.

Research-based literature in this area cautioned us against short-term GTA training programs (3, 26, 27). However, the reality of academic life is that sometimes long-term training programs in the specific area of teaching are not possible for a number of reasons (3, 28, 29), and this is where short term-training programs, even though not ideal, can be beneficial for GTA preparation (29, 30, 31). Moving forward, we would certainly advocate for use of these case studies in the context of an ongoing semester-long GTA training program. Each of the training cases can be discussed separately and incorporated into weekly laboratory preparation meetings. These would not be a major time commitment for the GTAs or the laboratory coordinator/instructor. Ideally, a follow-up classroom observation could be used to track adoption of pedagogy by the GTA in their lab sections to evaluate the effectiveness of the GTA trainings. As many of these GTAs will transition into the lifetime of teaching, this is worthy investment.

## SUPPLEMENTAL MATERIALS

These files are in Figshare repository DOI 10.6084/m9.figshare.7929020, https://figshare.com/s/21a8a7b50811dfd778f6.

**Appendix 1.** Training needs survey Likert-type questions and analysis

**Appendix 2.** Training needs survey open-ended questions and analysis

**Appendix 3.** Training cases, discussion questions and application exercises

**Appendix 4.** Lesson plan guide

**Appendix 5.** Knowledge Qualtrics survey

**Appendix 6.** Data Analysis Model Fit Details

**Appendix 7.** CIRTL Post-Survey questions and analysis

**Appendix 8**. “Live” CIRTL workshop data analysis.

## ACKNOWLEDGMENTS

We thank the Center for the Integration of Research, Teaching, and Learning (CIRTL) workshop participants for their participation in the Knowledge survey. We would also like to thank the graduate teaching assistants at Iowa State University, University of North Carolina Wilmington, and University of Georgia for their participation in the training needs survey and in training. This work was funded by National Science Foundation (NSF) Plant Genome Research Program (PGRP) grant #1444339. Any opinions, findings, and conclusions or recommendations expressed in this material are those of the authors and do not necessarily reflect the views of the NSF. Surveys and research were approved by the Institutional Review Board **(IRB ID 17-104**) at Iowa State University.

## Notes

**Conflicts of interest**: The authors declare that there are no conflicts of interest.

**Contributors and Supporting Agencies:** This work was partially supported by National Science Foundation (NSF) Plant Genome Research Program (PGRP) grant No. 1444339.

